# The *Drosophila* Gene Expression Tool (DGET) for expression analyses

**DOI:** 10.1101/075358

**Authors:** Yanhui Hu, Aram Comjean, Norbert Perrimon, Stephanie Mohr

**Author notes:** Corresponding author: Stephanie Mohr.

## Abstract

**Background:** Next-generation sequencing technologies have greatly increased our ability to identify gene expression levels, including at specific developmental stages and in specific tissues. Gene expression data can help researchers understand the diverse functions of genes and gene networks, as well as help in the design of specific and efficient functional studies, such as by helping researchers choose the most appropriate tissue for a study of a group of genes, or conversely, by limiting a long list of gene candidates to the subset that are normally expressed at a given stage or in a given tissue.

**Results:** We report a *Drosophila* Gene Expression Tool (DGET, www.flyrnai.org/tools/dget/web/), which stores and facilitates search of RNA-Seq based expression profiles available from the modENCODE consortium and other public data sets. Using DGET, researchers are able to look up gene expression profiles, filter results based on threshold expression values, and compare expression data across different developmental stages, tissues and treatments. In addition, at DGET a researcher can analyze tissue or stage-specific enrichment for an inputted list of genes (e.g. ‘hits’ from a screen) and search for additional genes with similar expression patterns. We performed a number of analyses to demonstrate the quality and robustness of the resource. In particular, we show that evolutionary conserved genes expressed at high or moderate levels in both fly and human tend to be expressed in similar tissues. Using DGET, we compared whole tissue profile and sub-region/cell-type specific datasets and estimated the potential cause of false positives in one dataset. We also demonstrated the usefulness of DGET for synexpression studies by querying genes with similar expression profile to the mesodermal master regulator Twist.

**Conclusion:** Altogether, DGET provides a flexible tool for expression data retrieval and analysis with short or long lists of *Drosophila* genes, which can help scientists to design stage- or tissue-specific *in vivo* studies and do other subsequent analyses.

## Background

The application of next-generation sequence technologies to RNA analysis has opened the door to relatively rapid, large-scale analyses of gene expression. ‘Standard’ RNA-seq analysis, for example, can provide a snapshot of gene expression in specific cell types or tissues (Wang, Gerstein et al. 2009), and related technologies such as Ribo-seq (Michel and Baranov 2013) provide more refined views, such as a snapshot of what genes are actively transcribed in a given cell or tissue. For *Drosophila*, efforts such as the modENCODE project (mod, Roy et al. 2010, Cherbas, Willingham et al. 2011, Graveley, Brooks et al. 2011, Boley, Wan et al. 2014) have provided a baseline overview of expression under standard laboratory conditions for various cultured cell types, developmental stages, and tissues, as well as treatment conditions. Moreover, studies such as those investigating expression in sub-regions of the fly gut (Marianes and Spradling 2013, Dutta, Dobson et al. 2015) are providing increasingly detailed views of the baseline expression levels of various genes in various tissues, cell types and sub-regions. Altogether, these RNAseq data resources provide helpful starting points for analysis of other gene lists.

Resources such as FlyBase (dos Santos, Schroeder et al. 2015) make it possible to quickly view modENCODE data for a given gene and make these data generally accessible to the community. The value of these data to the community can be further increased by facilitating search of lists of genes. For example, for gene lists originating from whole-animal or cultured cell studies, or for studies based on a list of orthologs of genes from another species, it can be very helpful to get a picture of what stages or tissues normally express those genes, as that will help focus stage- or tissue-specific *in vivo* studies and other subsequent analyses. We implemented DGET to help scientists retrieve modENCODE expression data in batch mode. DGET also hosts other relevant RNA-Seq datasets published in individual studies, such as profiles of specific sub-regions and cell types of the *Drosophila* gut (Marianes and Spradling 2013, Dutta, Dobson et al. 2015). Here, we describe DGET and perform a number of analyses to demonstrate the quality and robustness of the resource.

## Results and Discussion

### Database content and features of the user interface (UI)

The DGET database contains processed RNA-Seq data from the modENCODE consortium (mod, Roy et al. 2010, Cherbas, Willingham et al. 2011, Graveley, Brooks et al. 2011, Boley, Wan et al. 2014), as released by FlyBase (dos Santos, Schroeder et al. 2015), as well as other published datasets we obtained from supplemental tables (Marianes and Spradling 2013, Dutta, Dobson et al. 2015, Clough and Barrett 2016). The DGET UI has two tabs (Figure 1).

**Figure 1a.**
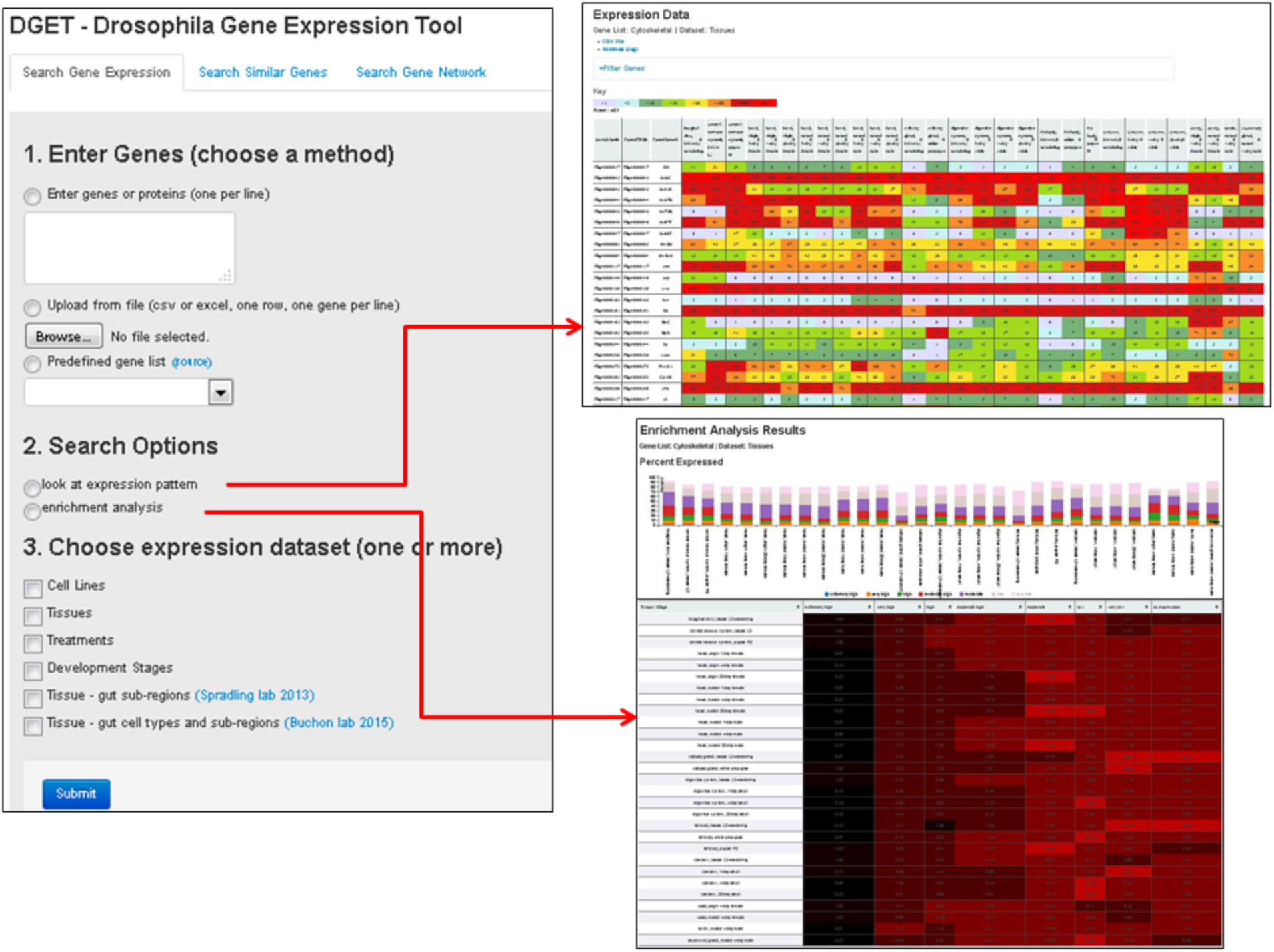
The DGET user interface. On the “Search Gene Expression” page, users can input a gene list by pasting *Drosophila* gene or protein symbols or IDs into the text box, or by uploading a file. The specific identifiers accepted are FlyBase Gene Identifier (FBgn), gene symbol, CG number, and full gene name. Users can choose to look at expression patterns or perform an enrichment analysis of the inputted list as compared with the underlying RNA-Seq data.

**Figure 1b.**
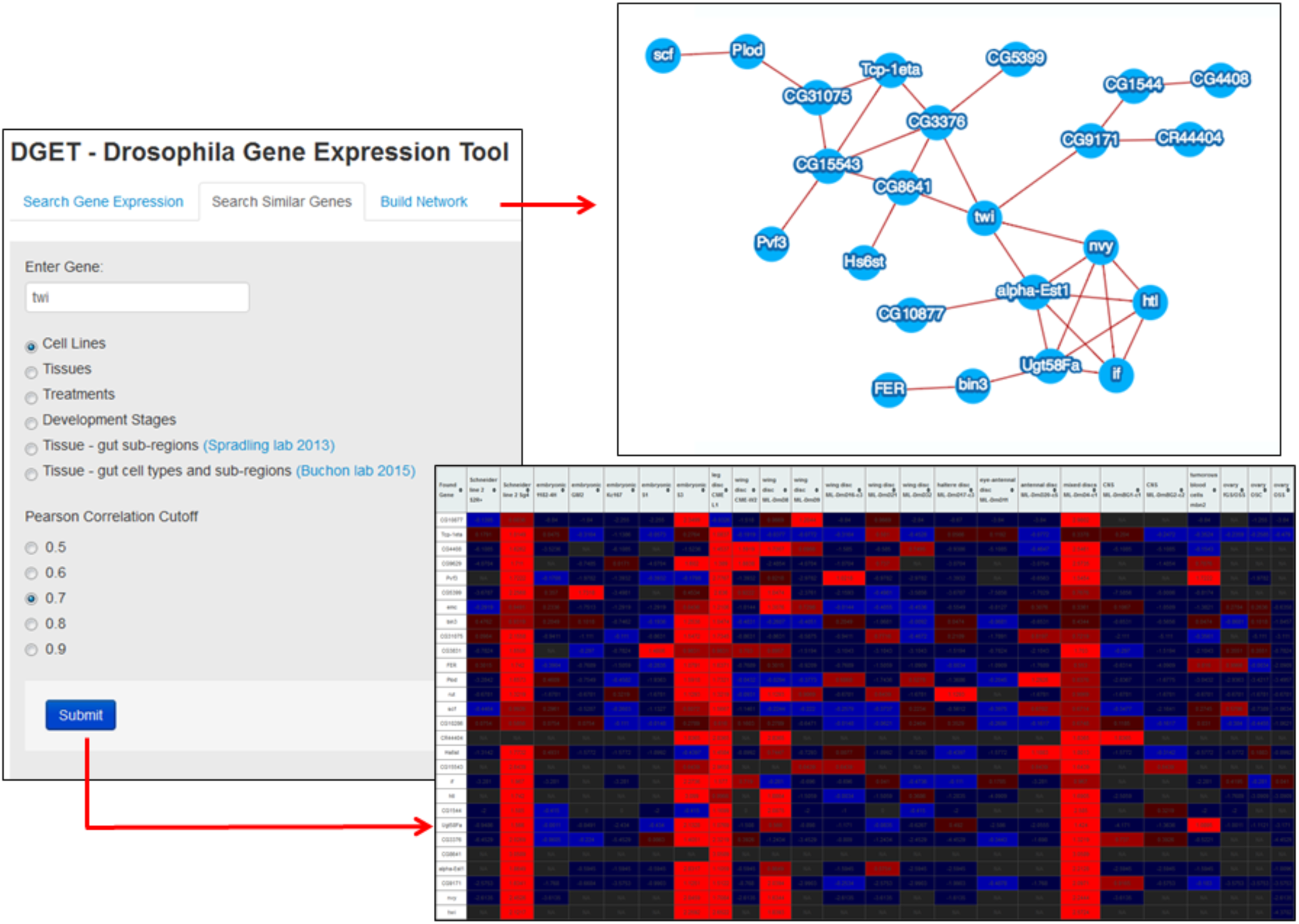
At the “Search Similar Genes” page, users can enter a gene symbol (or other accepted identifier) to find genes with similar expression patterns.

At the “Search Gene Expression” tab, users can enter a list of genes or choose one of the predefined gene classes from GLAD (Hu, Comjean et al. 2015), e.g. kinases, then specify the datasets to be displayed. There are two search options, “look at expression” and “enrichment analysis.” The results page for “look at expression” displays expression values in a heatmap format. At this results page, users have the option to download the relevant expression values; download the heatmap; and further filter the list by defining a cutoff, limit to specific dataset(s), or filtering out genes, for example with less than 1 RPKM value based on carcass and/or digestion system expression of 1 day adult. We used an RPKM cutoff of 1 because this is considered the cutoff for ‘no or extremely low expression’ at FlyBase. The results page for an enrichment analysis displays the distribution of genes at different expression levels using a bar graph and heatmap. The cutoff values for different levels are defined based on FlyBase guidelines.

Using the “Search Similar Genes” tab, users can enter a gene of interest and search for other genes with similar expression pattern based on Pearson correlation score. Users have the options to download the list of genes with similar expression patterns, a heatmap, and a normalized heatmap.

### Expression pattern of Drosophila regulatory genes

When genome-scale screening is not practical to do, a common approach is to select a specific subset of genes to start with, such as a group of genes with related activities. The most frequently screened sub-sets of genes are important regulatory genes including genes that encode kinases, phosphatases, transcription factors, or canonical signal transduction pathways components. Our expectation is that these regulatory genes, which appear to be re-used in many contexts, will be expressed in many tissues. To test this, we analyzed the expression patterns of several *Drosophila* regulatory gene classes defined by GLAD (Hu, Comjean et al. 2015). These included canonical signal transduction pathway genes, kinases, phosphatases, transcription factors, secreted proteins, and receptors. The percentages of expressed genes were calculated across all tissues profiled using a RPKM of 1 or above as a cutoff for expressed versus not expressed (Figure 2). About 70-90% of the genes categorized as encoding canonical signal transduction pathway components, kinases, phosphatases, or transcription factors are expressed in each of the major tissue categories profiled, whereas only 30-60% of receptor or secreted proteins are detected in any given tissue.

**Figure 2.**
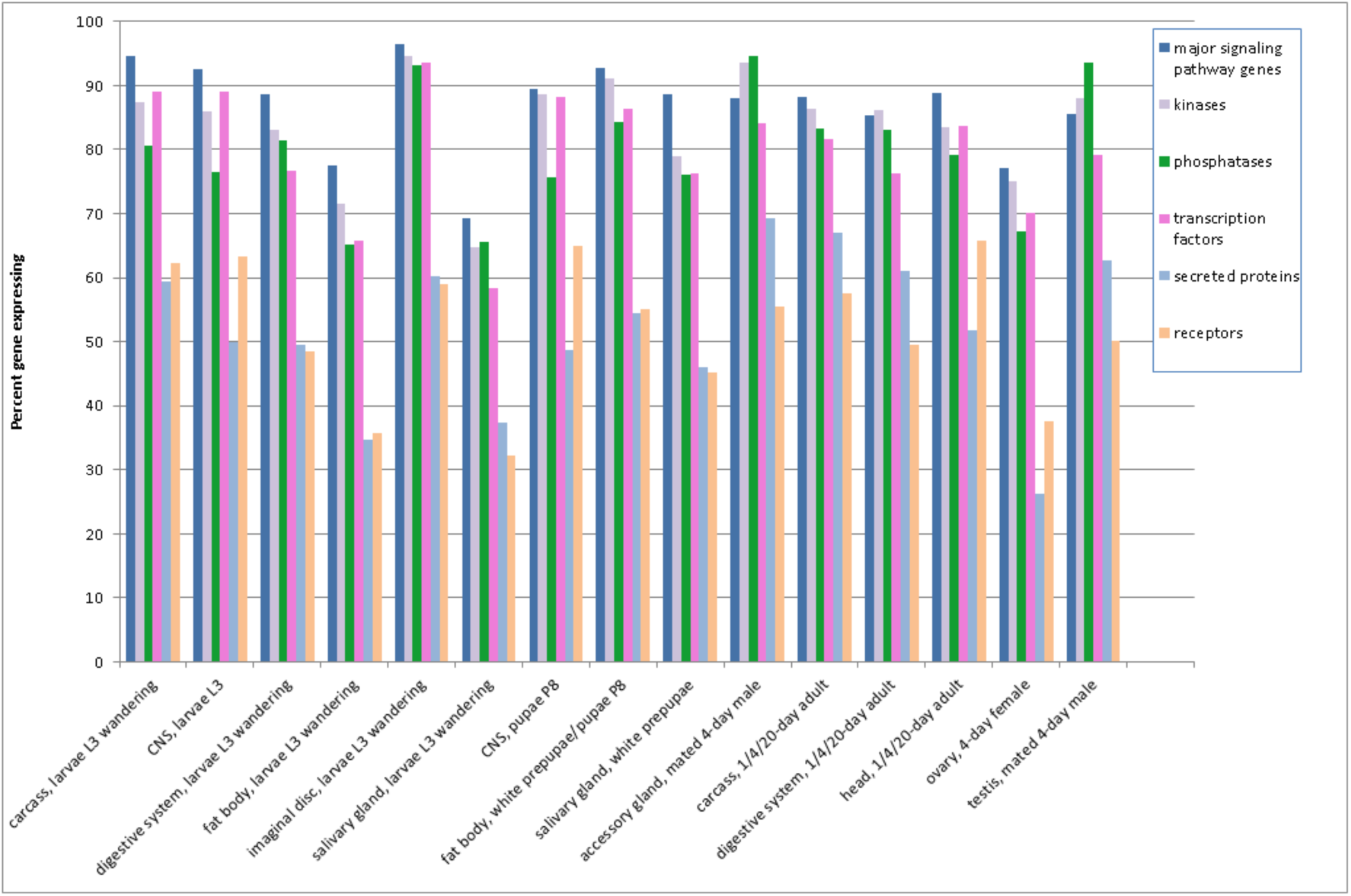
Expression patterns of genes in major Drosophila regulatory gene groups.

### Correlation of expression with confidence in an ortholog relationship

It is well established that the evolutionary conservation of proteins correlates with conservation at the level of biological and/or biochemical functions. *Drosophila* is a model organism of particular interest for which a wide variety of molecular genetic tools are readily available. Particularly, *Drosophila* models have been developed for a number of human diseases (Perrimon, Bonini et al. 2016). According to DIOPT, 9,705 of 13,902 protein-coding genes in *Drosophila* are predicted to have human ortholog(s) (Hu, Flockhart et al. 2011). Using DGET we analyzed the expression levels of the subset of *Drosophila* genes for which there is evidence that they are conserved in the human genome. Specifically, we analyzed subsets of genes scoring as putative human orthologs of fly genes at different levels of confidence, as defined by the DIOPT score (Hu, Flockhart et al. 2011). We found a strong correlation of percent expressed genes with DIOPT score (Figure 3). For example, for genes that have a high-confidence ortholog relationship (DIOPT score of 7 or above), almost all are expressed across all tissues. By contrast, for genes for which DIOPT analysis suggests that there is no evidence of a human ortholog (i.e. none of the 10 ortholog algorithms queried with DIOPT predict an ortholog), only 20-50% are expressed in each of the major tissue categories profiled. We suspect that this correlation is driven by essential genes, which are more conserved evolutionary. We also note that gene set enrichment for the set of high-confidence orthologs indicates that “kinases” and “nucleotide binding” among the top 20 enriched sets, indicating that the set of regulatory genes analyzed above has overlap with this set.

**Figure 3.**
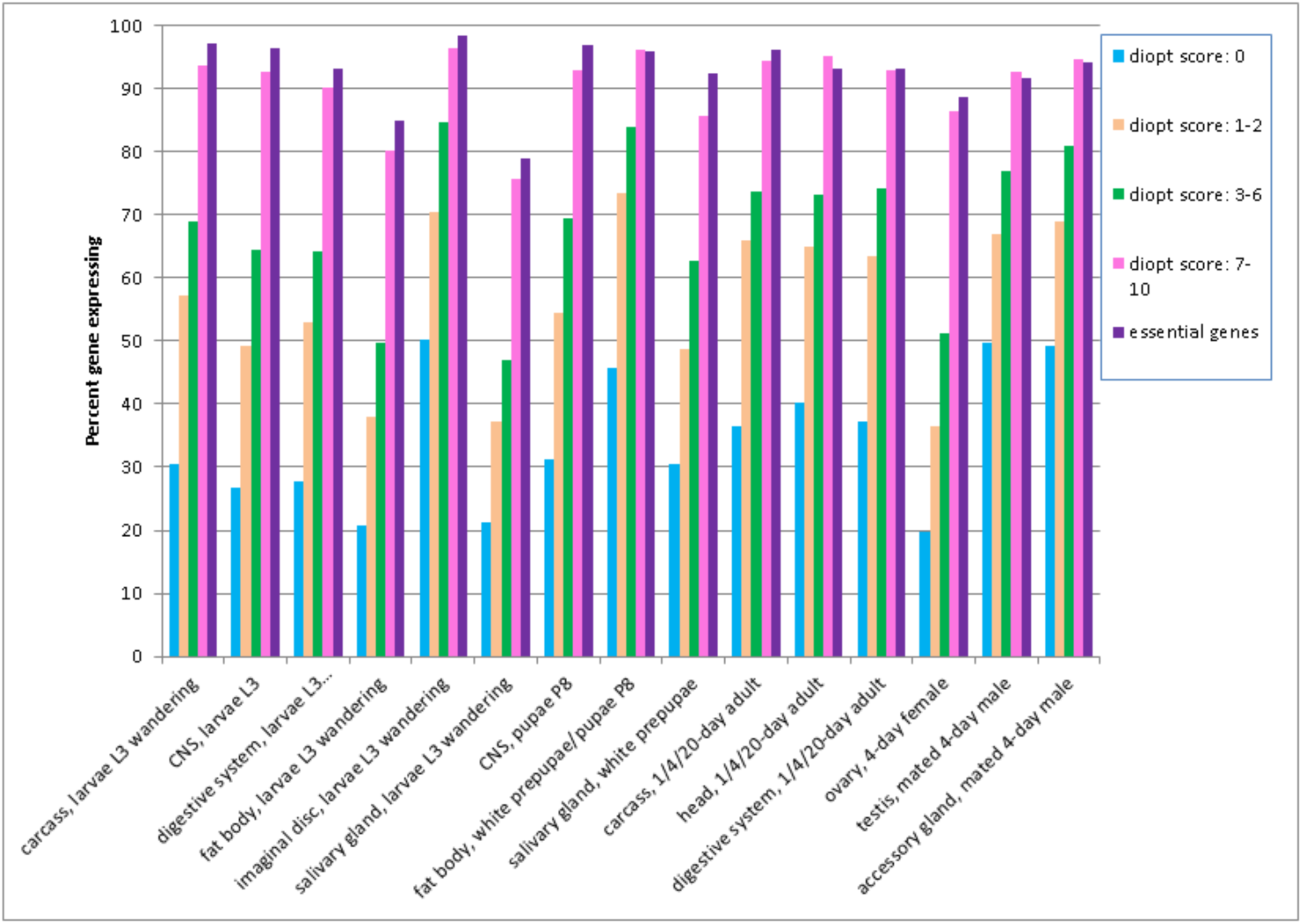
Relationship between expression levels and gene conservation. *Drosophila* genes that are conserved in the human genome at different confidence levels (i.e. different DIOPT scores) were analyzed by DGET. We found that across all tissues, expression levels correlate with confidence in the ortholog relationship. That is, in general, genes with higher DIOPT scores vs. human genes have higher expression levels. Genes with DIOPT scores of 7-10 (light purple bars) have similar expression patterns as compared with *Drosophila* essential genes (dark purple bars); i.e. in both cases, the genes are likely to be expressed in many or all tissues.

We next analyzed the 418 *Drosophila* essential genes identified by Spradling et al (Spradling, Stern et al. 1999) using a large-scale single P-element insertion fly stock collection. The proportions of essential genes expressed at detectable levels in various tissues are very similar to the genes with DIOPT score 7-10 (Figure 3, light purple and dark purple bars) with a Pearson correlation coefficient equal to 0.92.

### Expression patterns of *Drosophila* orthologs of human genes that are highly expressed in specific tissues

Next, we asked whether genes conserved between human and *Drosophila* are also expressed in similar patterns. We used the tissue-based human proteome annotation available at the Human Protein Atlas (HPA) (www.proteinatlas.org) (Uhlen, Fagerberg et al. 2015), as the source for tissue-specific expression, and retrieved the set of human genes that are expressed in specific tissues. Next, we mapped these human genes to *Drosophila* orthologs using DIOPT (Hu, Flockhart et al. 2011), filtering out low rank ortholog pairs (see Materials and Methods), and analyzed the expression patterns of these high-confidence orthologs in *Drosophila* tissues using DGET (Figure 4). The results of our analysis using all annotated proteins without a filter did not clearly demonstrate conservation of expression patterns. However, an analysis limited to genes expressed at high or moderate levels (as annotated by HPA) from high confident annotation (i.e. excluding HPA “reliability” value of “uncertain”), indicates that gene expression patterns are conserved in similar tissues in *Drosophila*. For example, as a group, orthologs of genes highly expressed in the human cerebellum, cerebral cortex, lateral ventricle or hippocampus are highly expressed in the *Drosophila* central nervous system (CNS) or head, at both larval and adult stages, and orthologs of genes highly expressed in human testis are also highly expressed in the *Drosophila* testis. Moreover, orthologs of genes from some organs of the human digestive system, such as stomach, duodenum or small intestine, are also highly expressed in the *Drosophila* digestive system. To further compare the expression patterns of genes expressed in the human and *Drosophila*, digestive systems, we analyzed the *Drosophila* gut sub-region data from Dutta et al. (Dutta, Dobson et al. 2015) (Figure 5). Orthologs of genes highly expressed in the human salivary gland and esophagus are highly expressed in the R1 upstream region, and orthologs of genes highly expressed in the human rectum, colon or appendix are more biased towards expression in the R5 downstream region. Fly orthologs of genes highly expressed in the human stomach, duodenum and small intestine are detected throughout the samples corresponding to R1 to R5.

**Figure 4.**
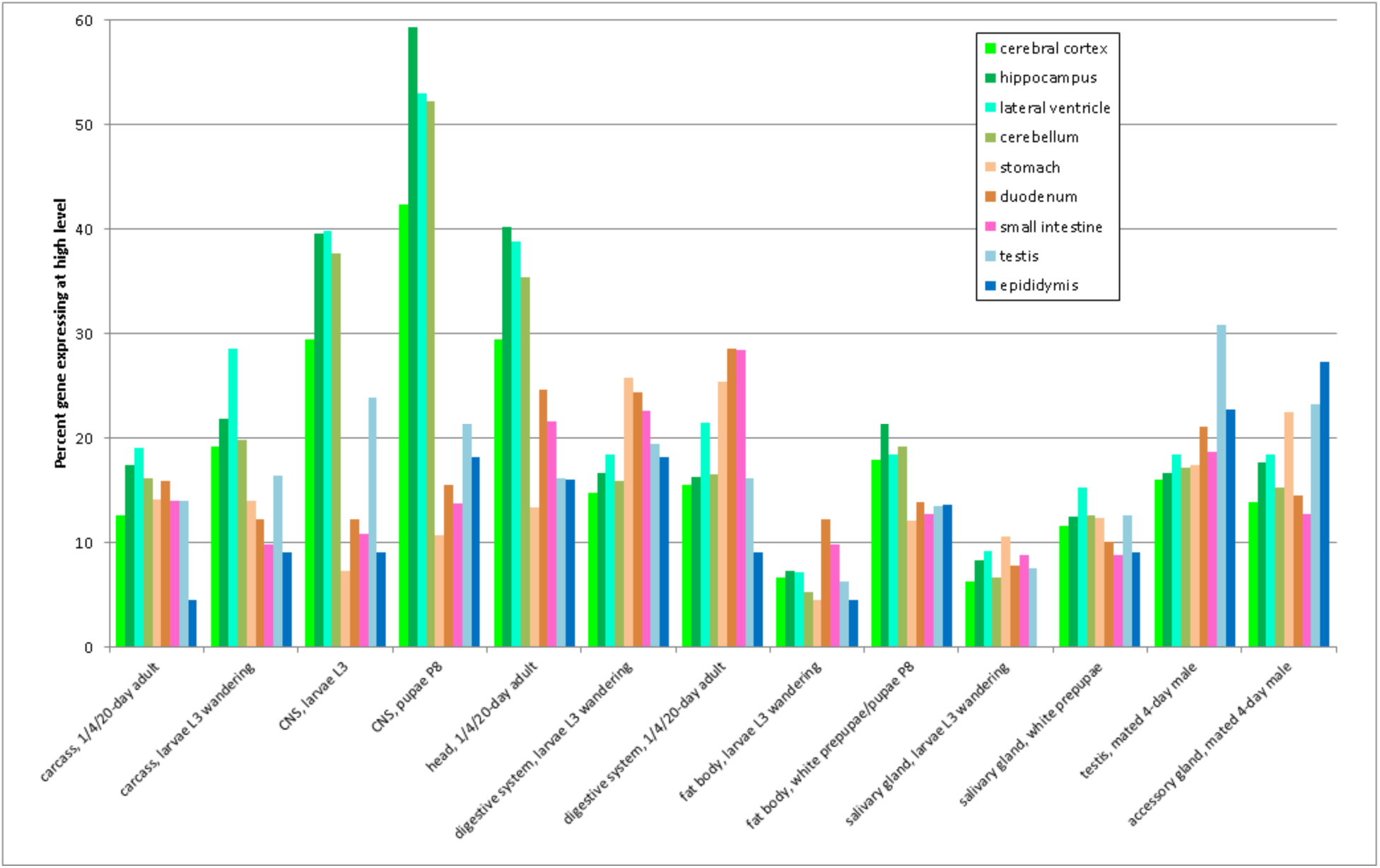
Comparison of gene expression patterns in humans and *Drosophila.* High-confidence *Drosophila* orthologs of genes that are highly expressed in the small intestine, ovary, testis, cerebellum, cerebral cortex, or other tissues were analyzed using DGET. For at least some tissues, we see a correlation between genes highly expressed in specific human tissues (e.g. cerebellum, testis) and the expression of orthologs in cognate tissue sample(s) available for *Drosophila* (e.g. CNS or head, testis).

**Figure 5.**
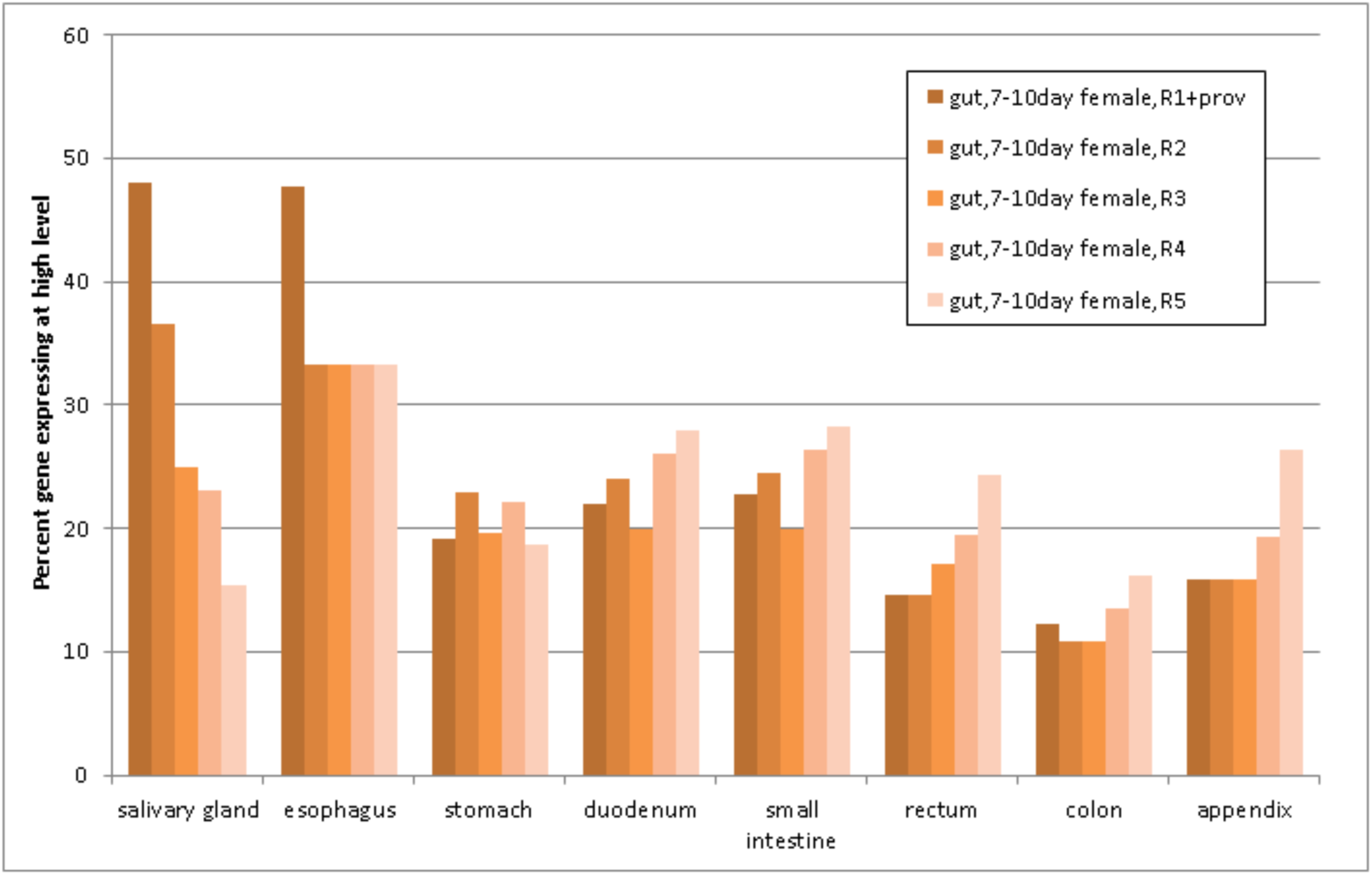
Comparison of *Drosophila* gut sub-region data with the human digestive system.

### Mining information from distinct but related fly gut gene expression data sets

We next sought to compare the results of whole-gut profiling with results from profiling of specific sub-regions or cell types with the goal of identifying genes only expressed in specific sub-populations. Our rationale for the analysis was to determine the likelihood that genes expressed in a sub-population are missed in expression analysis of an entire organ. This type of false negative analysis should provide helpful information for interpreting results of whole-organ or whole-tissue studies. Thus, we compared the whole gut profiling data obtained by modENCODE consortium for 20 day old adult flies (mod, Roy et al. 2010) with data generated by profiling sub-regions of the midgut in 16-20 day old adult flies (Marianes and Spradling 2013). Whole gut profiling indicates that 9,109 genes are expressed in the gut of 20 day old adult flies (RPKM cutoff value of 1). Among the 4,790 protein-coding genes not detected as expressed in the whole-gut study, 136 genes are detected in at least 3 sub-regions of the gut (RPKM>=3). These genes are either false negative in whole gut profiling or false positive in sub-region profiling. We next did a gene set enrichment analysis with these 136 genes and found that stress response genes, such as heat-shock genes (*Hsp70Aa, Hsp70Ab, Hsp70Ba, Hsp70Bbb*) are enriched (P value= 3.05E-07). This suggests that the sample used for sub-region profiling was associated with some level of stress. Comparing the list of 136 genes with the *Drosophila* essential gene list, we found only one overlapping gene. In addition, only 23 of the 136 genes have DIOPT score 7-10 when mapping to human genes. Thus, small fraction of these genes might be the false negative with whole tissue profiling while majority of the genes are likely to be the false positives not normally present in the gut under non-stress conditions.

### Synexpression analysis for transcription factor Twist

Expression profiling is a powerful approach to identify functionally related genes, as genes showing synexpression often operate in similar pathways and/or processes (see for example (Dequeant, Fagegaltier et al. 2015)). We tested DGET for its usefulness for synexpression studies by querying genes with similar expression profile to the mesodermal master regulator Twist. DGET preferentially retrieved Twist target genes with cell line data as well as development data. For example, among the top 27 genes that share similar expression with Twist in cell lines (Pearson correlation co-efficiency cut off = 0.7), 11 of them are Twist target genes based on Chip-on-chip data (Sandmann, Girardot et al. 2007), and 8 of the 11 genes are high-confidence (Table 1). The enrichment p-value for Twist target genes is 8.70E-04 and 3.05E-05 for high-confidence targets. We observed a less significant enrichment with development data (p-value 5.00E-02 for all Twist target genes and p-value of 2.70E-03 for high-confidence targets), likely reflecting the diversity of cell types present in the developmental data and that not enough cells express *twist*. Thus, DGET will be very powerful when applied to RNA-seq data sets from single cell or groups of homogeneous cell populations.

**Table 1.** DGET similar gene search results for Twist with cell line data

| FBgn | Gene | Correlation score | Twist target?* |
| --- | --- | --- | --- |
| FBgn0005636 | nvy | 0.910987 | Yes, high confident |
| FBgn0031738 | CG9171 | 0.88094 | Yes, high confident |
| FBgn0015568 | alpha-Est1 | 0.831094 |  |
| FBgn0035733 | CG8641 | 0.816603 |  |
| FBgn0034997 | CG3376 | 0.813417 | Yes, low confident |
| FBgn0040091 | Ugt58Fa | 0.799835 |  |
| FBgn0039827 | CG1544 | 0.773761 |  |
| FBgn0010389 | htl | 0.772568 | Yes, high confident |
| FBgn0001250 | if | 0.769353 | Yes, high confident |
| FBgn0039799 | CG15543 | 0.765649 |  |
| FBgn0038755 | Hs6st | 0.76281 | Yes, high confident |
| FBgn0265577 | CR44404 | 0.76095 |  |
| FBgn0037439 | CG10286 | 0.745739 |  |
| FBgn0025682 | scf | 0.744414 |  |
| FBgn0003301 | rut | 0.74375 | Yes, low confident |
| FBgn0036147 | Plod | 0.73896 |  |
| FBgn0000723 | FER | 0.738927 | Yes, low confident |
| FBgn0034804 | CG3831 | 0.735346 |  |
| FBgn0051075 | CG31075 | 0.731916 |  |
| FBgn0263144 | bin3 | 0.72961 | Yes, high confident |
| FBgn0000575 | emc | 0.728894 | Yes, high confident |
| FBgn0038353 | CG5399 | 0.724139 |  |
| FBgn0085407 | Pvf3 | 0.720044 | Yes, high confident |
| FBgn0036857 | CG9629 | 0.716929 |  |
| FBgn0039073 | CG4408 | 0.714359 |  |
| FBgn0037632 | Tcp-1eta | 0.702547 |  |
| FBgn0038804 | CG10877 | 0.701509 |  |

\* Twist targets as defined in (Sandmann, Girardot et al. 2007)

## Concluding Remarks

In summary, DGET makes it possible to retrieve and compare *Drosophila* gene expression patterns generated by various groups using RNA-Seq. The tool can help scientists design experiments based on expression and analyze experiment results. The backend database for DGET is designed to easily accommodate the addition of new high quality RNA-Seq datasets as they become available. Finally, although the anatomy of human and *Drosophila* are quite different, by using DGET, we demonstrate that expression patterns of genes that are conserved and highly expressed are conserved between human and *Drosophila* in many matching tissues, underscoring the utility of the *Drosophila* model to understand the role of human genes with unknown functions.

## Methods

### Data retrieval

Processed modENCODE data were retrieved from FlyBase (ftp://ftp.flybase.net/releases/current/precomputed_files/genes/gene_rpkm_report_fb_2015_05.tsv.gz). Data published by Marianes and Spradling (Marianes and Spradling 2013) were retrieved from NCBI Gene Expression Omnibus at (http://www.ncbi.nlm.nih.gov/geo/query/acc.cgi?acc=GSE47780). Data published by Dutta et al (Dutta, Dobson et al. 2015) were retrieved from the flygut-seq website (http://flygutseq.buchonlab.com/resources). Data retrieved were mapped to FlyBase identifiers from release 2015_5 and formatted for upload into the FlyRNAi database (Hu, Flockhart et al. 2011).

### Expression pattern analysis

Human protein expression data were retrieved from proteinatlas.org and tissue-specific genes were selected using the file “ProteinAtlas_Normal_tissue_vs14.” Proteins with high or medium expression levels with a reliability value of “supportive” were selected. Proteins expressed in a broad range of tissues (i.e. more than 5 tissues) were filtered out. DIOPT vs5 was used to map genes from human to *Drosophila (Hu, Flockhart et al. 2011)*. ‘Ortholog pair rank’ was added at recent DIOPT release 5.2.1 (http://www.flyrnai.org/DRSC-ORH.html#versions).*Drosophila* genes with high or moderate rank were selected. The high/moderate rank mapping include the gene pairs that are best score in either forward or reverse mapping (and DIOPT score >1) as well as gene pairs with DIOPT score >3 if not best score either way.

### Implementation

DGET was implemented using php and JavaScript with MySQL database for data store. It is hosted on a server provided by the Research IT Group (RITG) at Harvard Medical School. The MySQL database is also hosted on a server provided by RITG. Plotting of heat-maps for svg download is done in R using the gplot heatmap function. Website bar charts are drawn using the 3C.js plotting package. The php symfony framework scaffold is used to create DGET webpages and forms.

## Declarations

### Authors’ contributions

YH designed and tested the application, implemented the back-end of the application, performed the analysis and drafted the manuscript. AC implemented the user interface and contributed to the back-end of the application. NP provided critical input on key features and the analysis as well as edited the manuscript. SEM provided oversight and critical input on key features and the analysis, and helped draft the manuscript. All authors read and approved the final manuscript.

## References

Boley, N., K. H. Wan, P. J. Bickel and S. E. Celniker (2014). "Navigating and mining modENCODE data." Methods 68(1): 38–47.

Cherbas, L., A. Willingham, D. Zhang, L. Yang, Y. Zou, B. D. Eads, J. W. Carlson, J. M. Landolin, P. Kapranov, J. Dumais, A. Samsonova, J. H. Choi, J. Roberts, C. A. Davis, H. Tang, M. J. van Baren, S. Ghosh, A. Dobin, K. Bell, W. Lin, L. Langton, M. O. Duff, A. E. Tenney, C. Zaleski, M. R. Brent, R. A. Hoskins, T. C. Kaufman, J. Andrews, B. R. Graveley, N. Perrimon, S. E. Celniker, T. R. Gingeras and P. Cherbas (2011). “The transcriptional diversity of 25 Drosophila cell lines.” Genome Res 21(2): 301–314.

Clough, E. and T. Barrett (2016). “The Gene Expression Omnibus Database.” Methods Mol Biol 1418: 93–110.

Dequeant, M. L., D. Fagegaltier, Y. Hu, K. Spirohn, A. Simcox, G. J. Hannon and N. Perrimon (2015). “Discovery of progenitor cell signatures by time-series synexpression analysis during Drosophila embryonic cell immortalization.” Proc Natl Acad Sci U S A 112(42): 12974–12979.

dos Santos, G., A. J. Schroeder, J. L. Goodman, V. B. Strelets, M. A. Crosby, J. Thurmond, D. B. Emmert, W. M. Gelbart and C. FlyBase (2015). “FlyBase: introduction of the Drosophila melanogaster Release 6 reference genome assembly and large-scale migration of genome annotations.” Nucleic Acids Res 43(Database issue): D690–697.

Dutta, D., A. J. Dobson, P. L. Houtz, C. Glasser, J. Revah, J. Korzelius, P. H. Patel, B. A. Edgar and N. Buchon (2015). “Regional Cell-Specific Transcriptome Mapping Reveals Regulatory Complexity in the Adult Drosophila Midgut.” Cell Rep 12(2): 346–358.

Graveley, B. R., A. N. Brooks, J. W. Carlson, M. O. Duff, J. M. Landolin, L. Yang, C. G. Artieri, M. J. van Baren, N. Boley, B. W. Booth, J. B. Brown, L. Cherbas, C. A. Davis, A. Dobin, R. Li, W. Lin, J. H. Malone, N. R. Mattiuzzo, D. Miller, D. Sturgill, B. B. Tuch, C. Zaleski, D. Zhang, M. Blanchette, S. Dudoit, B. Eads, R. E. Green, A. Hammonds, L. Jiang, P. Kapranov, L. Langton, N. Perrimon, J. E. Sandler, K. H. Wan, A. Willingham, Y. Zhang, Y. Zou, J. Andrews, P. J. Bickel, S. E. Brenner, M. R. Brent, P. Cherbas, T. R. Gingeras, R. A. Hoskins, T. C. Kaufman, B. Oliver and S. E. Celniker (2011). “The developmental transcriptome of Drosophila melanogaster.” Nature 471(7339): 473–479.

Hu, Y., A. Comjean, L. A. Perkins, N. Perrimon and S. E. Mohr (2015). “GLAD: an Online Database of Gene List Annotation for Drosophila.” J Genomics 3: 75–81.

Hu, Y., I. Flockhart, A. Vinayagam, C. Bergwitz, B. Berger, N. Perrimon and S. E. Mohr (2011). “An integrative approach to ortholog prediction for disease-focused and other functional studies.” BMC Bioinformatics 12: 357.

Marianes, A. and A. C. Spradling (2013). “Physiological and stem cell compartmentalization within the Drosophila midgut.” Elife 2: e00886.

Michel, A. M. and P. V. Baranov (2013). “Ribosome profiling: a Hi-Def monitor for protein synthesis at the genome-wide scale.” Wiley Interdiscip Rev RNA 4(5): 473–490.

mod, E. C., S. Roy, J. Ernst, P. V. Kharchenko, P. Kheradpour, N. Negre, M. L. Eaton, J. M. Landolin, C. A. Bristow, L. Ma, M. F. Lin, S. Washietl, B. I. Arshinoff, F. Ay, P. E. Meyer, N. Robine, N. L. Washington, L. Di Stefano, E. Berezikov, C. D. Brown, R. Candeias, J. W. Carlson, A. Carr, I. Jungreis, D. Marbach, R. Sealfon, M. Y. Tolstorukov, S. Will, A. A. Alekseyenko, C. Artieri, B. W. Booth, A. N. Brooks, Q. Dai, C. A. Davis, M. O. Duff, X. Feng, A. A. Gorchakov, T. Gu, J. G. Henikoff, P. Kapranov, R. Li, H. K. MacAlpine, J. Malone, A. Minoda, J. Nordman, K. Okamura, M. Perry, S. K. Powell, N. C. Riddle, A. Sakai, A. Samsonova, J. E. Sandler, Y. B. Schwartz, N. Sher, R. Spokony, D. Sturgill, M. van Baren, K. H. Wan, L. Yang, C. Yu, E. Feingold, P. Good, M. Guyer, R. Lowdon, K. Ahmad, J. Andrews, B. Berger, S. E. Brenner, M. R. Brent, L. Cherbas, S. C. Elgin, T. R. Gingeras, R. Grossman, R. A. Hoskins, T. C. Kaufman, W. Kent, M. I. Kuroda, T. Orr-Weaver, N. Perrimon, V. Pirrotta, J. W. Posakony, B. Ren, S. Russell, P. Cherbas, B. R. Graveley, S. Lewis, G. Micklem, B. Oliver, P. J. Park, S. E. Celniker, S. Henikoff, G. H. Karpen, E. C. Lai, D. M. MacAlpine, L. D. Stein, K. P. White and M. Kellis (2010). “Identification of functional elements and regulatory circuits by Drosophila modENCODE.” Science 330(6012): 1787–1797.

Perrimon, N., N. M. Bonini and P. Dhillon (2016). “Fruit flies on the front line: the translational impact of Drosophila.” Dis Model Mech 9(3): 229–231.

Sandmann, T., C. Girardot, M. Brehme, W. Tongprasit, V. Stolc and E. E. Furlong (2007). “A core transcriptional network for early mesoderm development in Drosophila melanogaster.” Genes Dev 21(4): 436–449.

Spradling, A. C., D. Stern, A. Beaton, E. J. Rhem, T. Laverty, N. Mozden, S. Misra and G. M. Rubin (1999). “The Berkeley Drosophila Genome Project gene disruption project: Single P-element insertions mutating 25% of vital Drosophila genes.” Genetics 153(1): 135–177.

Uhlen, M., L. Fagerberg, B. M. Hallstrom, C. Lindskog, P. Oksvold, A. Mardinoglu, A. Sivertsson, C. Kampf, E. Sjostedt, A. Asplund, I. Olsson, K. Edlund, E. Lundberg, S. Navani, C. A. Szigyarto, J. Odeberg, D. Djureinovic, J. O. Takanen, S. Hober, T. Alm, P. H. Edqvist, H. Berling, H. Tegel, J. Mulder, J. Rockberg, P. Nilsson, J. M. Schwenk, M. Hamsten, K. von Feilitzen, M. Forsberg, L. Persson, F. Johansson, M. Zwahlen, G. von Heijne, J. Nielsen and F. Ponten (2015). “Proteomics. Tissue-based map of the human proteome.” Science 347(6220): 1260419.

Wang, Z., M. Gerstein and M. Snyder (2009). “RNA-Seq: a revolutionary tool for transcriptomics.” Nat Rev Genet 10(1): 57–63.

